# Error-related cognitive control and behavioural adaptation mechanisms in the context of motor functioning and anxiety

**DOI:** 10.1101/2020.09.04.281758

**Authors:** Marta Topor, Bertram Opitz, Hayley C. Leonard

**Affiliations:** School of Psychology, University of Surrey, Guildford, UK

**Keywords:** Motor skills, cognitive control, behavioural adaptation, anxiety, error-related negativity, post-error slowing

## Abstract

Previous research suggests that there is an interaction between cognitive and motor processes. This has been investigated throughout development and in different conditions related to motor impairment. The current study addressed a gap in the literature by investigating this interaction in the general population of healthy adults with different profiles of motor proficiency by focusing on error-related cognitive control and behavioural adaptation mechanisms. In addition, the impact of these processes was assessed in terms of experienced anxiety. Forty healthy adults were divided into high and low motor proficiency groups based on an assessment of their motor skills. Using electroencephalography (EEG) during a flanker task, error-related negativity (ERN) was measured as the neural indicator of cognitive control. Post-error slowing (PES) was measured to represent behavioural adaptation. Participants also completed an anxiety assessment questionnaire. Participants in the high motor proficiency group achieved better task accuracy and showed relatively enhanced cognitive control through increased ERN. Contrastingly, individuals in the lower motor proficiency group achieved poorer accuracy whilst showing some evidence of compensation through increased PES. Anxiety was found to be associated with motor functioning, but the study could not provide evidence that this was related to cognitive or behavioural control mechanisms. The interaction between cognitive and motor processes observed in this study is unique for healthy and sub-clinical populations and provides a baseline for the interpretation of similar investigations in individuals with motor disorders.

## 1. Introduction

The relationship between motor and cognitive functioning (from now referred to as the M-C interaction) is an important topic explored through different research approaches. It has been studied specifically in the context of child development (van der Fels et al., 2015) and health conditions affecting movement including, amongst others, neurodevelopmental conditions such as developmental coordination disorder (DCD; Sartori, Valentini, & Fonseca, 2020), neurodegenerative conditions such as Parkinson’s disease (PD; Sollinger, Goldstein, Lah, Levey, & Factor, 2010), as well as stroke (Plummer et al., 2013), and acquired brain injury (Damiano, Zampieri, Ge, Acevedo, & Dsurney, 2016). Research concerning the M-C interaction in the adult general population is rare but has recently gained more attention with the aim of investigating how executive and cognitive processes may be associated with motor ability and proficiency (Ludyga, Pühse, Gerber, & Herrmann, 2019; Marchetti et al., 2015; Stuhr, Hughes, & Stöckel, 2018).

### 1.1. Observations of the motor-cognitive interaction

The latest advancements in the attempt to understand the M-C interaction in the general population suggest that this interaction may link specific faculties of executive and cognitive functioning with different motor domains (Ludyga et al., 2019; Marchetti et al., 2015). For instance, reports suggest that working memory is associated with fine motor skills (Stuhr et al., 2018) as well as locomotor skills (Ludyga et al., 2019). On the other hand, inhibitory control may be associated with balance (Rigoli, Piek, Kane, & Oosterlan, 2012; Stuhr et al., 2018). A common observation across these studies suggests that the specific cognitive processes may help to facilitate control over an individual’s action in order to improve task performance. For instance, Stuhr et al. (2018) argue that the M-C interaction is mediated by task difficulty, and cognitive control processes are engaged to facilitate a successful completion of tasks that are difficult or novel. Marchetti et al. (2015), on the other hand, explore the possibility that the engagement of cognitive processes such as inhibition may facilitate good performance on motor tasks that are challenging and require a strategic approach. Thus, the M-C interaction can be viewed to be goal-directed, helping individuals to perform to the best of their ability despite the challenges of the tasks and activities being undertaken (Marchetti et al., 2015; Stuhr et al., 2018). Stuhr et al. (2018) suggest that this could be the explanation for why most of the available empirical evidence for the M-C interaction comes from populations with impaired motor ability. Tasks which require motor responses are more challenging for such individuals and thus differences in the M-C interaction process may be observed which, in turn, inform us about the nature of this relationship.

It is however important to also study the M-C interaction in the general population with no motor impairments. It will aid the understanding of the general patterns and risk factors associated with cognitive and motor functioning interaction. Different levels of motor proficiency can be observed among healthy individuals who, albeit to a lesser extent, could also be affected by motor and cognitive functioning challenges similar to those experienced by people with motor disorders. The understanding of the general patterns of the M-C interaction will also aid the interpretation of already existing research and direct the development of future projects. For instance, if the pattern of M-C interactions in individuals with motor disorders is observed to be different to that in the general population, it would indicate syndrome-specific changes which may occur throughout development. This may not be understood until the general patterns of the M-C interaction in healthy individuals is established.

The recent studies by Ludyga et al. (2019), Marchetti et al. (2015) and Stuhr et al. (2018) are very informative regarding the nature of the M-C interaction in the general population but rely on cognitive-behavioural approaches. In all of these studies, participants’ motor skills were tested using standardised motor assessments whilst cognitive functioning was tested using a variety of cognitive tasks. However, the M-C interaction could be studied in more depth by investigating control process at the neural and cognitive in addition to behavioural levels therefore, the current study aims to apply such an approach.

### 1.2. Control processes on the neural level

To the best of our knowledge, the M-C interaction on the neural level has only been studied by observing the presentation and performance of individuals with motor disorders. One example is of developmental coordination disorder (DCD). In DCD, individuals experience significant difficulties with the execution of coordinated movements but also problems with executive functions (Leonard, Bernardi, Hill, & Henry, 2015; Sartori et al., 2020). Querne et al. (2008) conducted a study using functional magnetic resonance imagery (fMRI) with children completing a standard Go/NoGo paradigm, in which responses must be provided or inhibited depending on the instructions. Children with DCD achieved similar accuracy to healthy children on the task, despite slower responses. The fMRI results suggest that children with DCD may have a stronger activation of the anterior cingulate cortex (ACC), which may help to maintain good accuracy on the task whilst compensating for motor difficulties.

The ACC is a key structure within the fronto-striatal network. It is the central part of the Human Error Processing Model proposed by Holroyd and Coles (2002), which has been updated over the years and recently resulted in a new comprehensive model suggested by Brown and Alexander (2017). The model presents a complex infrastructure of ACC activity which encompasses the processes of learning and updating of goals, available actions and their outcomes, as well as the selection and the execution of appropriate actions. A vital element of the action control models of the ACC is the prediction error (PE), which compares the expected outcomes to actual outcomes and reflects surprise, which may be positive or negative. The ACC uses the PE signal to evaluate the available actions and signals a mismatch between expected and actual outcomes which helps to navigate the selection of appropriate actions in the future (Robbins, 2009). PE can be measured in the form of an error-related negativity (ERN), a response-locked component of the event-related potential observed with electroencephalography (EEG) following error commission. It occurs about 50ms post erroneous response and larger amplitudes indicate stronger engagement of the ACC (Hauser et al., 2014; Shenhav, Botvinick, & Cohen, 2013). Although the ACC has been implicated in M-C interaction in DCD using fMRI, the ERN has not yet been studied in this population. However, it has recently been investigated in another neurodevelopmental motor disorder, Tourette’s syndrome (TS), which is characterised by difficulties with the control of involuntary movements (Bloch & Leckman, 2009). Schüller et al. (2018) showed that adult participants with TS could perform the stop-signal task, which requires the inhibition of responses when instructed to do so, with accuracy similar to that of healthy participants. However, their ERN amplitude was significantly larger. This pattern can be compared to the results observed using fMRI in children with DCD by Querne et al. (2008). Both studies suggest that participants with motor disorders perform cognitive tasks (Go/NoGo or stop-signal) with the same accuracy as their peers with no motor conditions but they engage additional cognitive efforts on the neural level. These enhanced cognitive processes may facilitate task performance of individuals with motor disorders as a way of compensation for their motor control difficulties. This highlights the importance of investigating both neural and behavioural patterns of performance control which is the aim of the current study.

### 1.3 Control processes on the behavioural level

Altered cognitive control processes reflecting compensation to aid task performance can be seen at the neural level with greater ACC activity (Querne et al., 2008; Schüller et al., 2018). These control processes can also be studied by how they change individual behaviour into strategies helping to achieve better accuracy and performance. These strategies can be understood as adaptations taken when the ACC detects a discrepancy between the expected and actual outcomes of an action (Brown & Alexander, 2017). One behavioural strategy that has been studied in this context is the post-error slowing (PES). PES is commonly investigated alongside the ERN and recent studies suggest that the magnitude of PES can be predicted by the neural correlates of the ERN signal (Chang, Chen, Li, & Li, 2014; Fu et al., 2019). The relationship between the ERN and PES has been found to persist throughout development and is associated with task accuracy (Ladouceur, Dahl, & Carter, 2007). The PES has not yet been investigated in motor disorders or in the context of motor ability in the general population. Such research is important because it may provide important insights into how enhanced cognitive control processes in the ACC may affect behavioural control, which can compensate for poor motor skills.

### 1.4. Control processes and anxiety

Whilst increased motor-cognitive interaction and cognitive control may be a compensation that leads to better performance, this increased control may also have implications for the individual’s emotional functioning, such as the experience of worries and anxiety. Emotional difficulties are commonly observed in those with poor motor skills, especially in clinical populations such as DCD or TS, who report high levels of anxiety (Bloch & Leckman, 2009; Hill & Brown, 2013). However, a recent study reported that even healthy adults with relatively poorer motor proficiency experience higher levels of anxiety than those with better motor skills (Rigoli et al., 2017). The source of these emotional difficulties is yet to be investigated although, so far it has been attributed to the environmental stress hypothesis (ESH; Cairney, Rigoli, & Piek, 2013) both in the context of motor disorders and poor motor ability in healthy individuals (Rigoli et al., 2017). The ESH was developed based on the experiences of children with DCD and suggests that motor difficulties expose individuals to negative experiences (such as not getting on with peers in school) that lead to negative self-perceptions and internalising problems. Thus, the current explanation for anxious tendencies in relation to motor difficulties focuses on individual experiences and external factors whilst the possible impact of cognitive processes is yet to be considered.

The application of compensatory processes to support task performance could be another possible explanation for the observed high levels of anxiety in individuals with poor motor skills. It has been suggested that individuals who engage enhanced cognitive control efforts to facilitate their performance and avoid errors may be more sensitive to error making and they may be characterised by increased worries when committing errors (Frank, Woroch, & Curran, 2005; Holroyd & Umemoto, 2016). In fact, enhanced ERN is commonly observed in individuals with anxiety disorders (Weinberg, Kotov, & Proudfit, 2015), as well as undiagnosed individuals with high anxiety, which indicates that enhanced error control may be associated with trait anxiety (Hajcak, McDonald, & Simons, 2003). Investigating the anxious tendencies of individuals with poor motor functioning through this lens would help to understand the emotional risk factors that may be associated with their motor difficulties. Linking these risk factors with functioning at behavioural, cognitive and neural levels will be an important step forward in understanding the M-C interaction and its consequences, and it is to this end that the current study was conducted.

### 1.5 The current study

The aim of the current study was to investigate the M-C interaction and its association with anxiety and worry in an adult general population with different motor proficiency profiles. By doing so, the study will elucidate whether the patterns and results seen in motor disorders are also present in the general population. Thus, the study applied methods similar to those previously used in research on DCD and TS but this time in two groups of healthy adults with higher motor proficiency (HMP group) and lower motor proficiency (LMP group). The patterns of cognitive and behavioural control mechanisms were measured through the ERN and PES during a standard Flanker task. In this task participants were instructed to respond to a string of arrows presented on the screen. The direction of the middle arrow indicated which button should be pressed. The task had three conditions which depended on the direction of the remaining arrows – congruent (same as the middle arrow), incongruent (opposite to the middle arrow) or neutral (the letter “v” was used instead of arrows).

Given that cognitive-behavioural task performance is often similar between those with and without motor disorders, it was expected that both the LMP and HMP groups would show a typical flanker effect, i.e. accuracy would be lower and reaction times higher for the incongruent trials compared to congruent and neutral conditions (Hypothesis 1). Considering the enhanced ACC activity in individuals with developmental motor disorders, it was expected that the LMP group would have larger ERN amplitudes and longer PES (Hypothesis 2). Although PES has not yet been studied in association with motor functioning, this prediction was based on the close relationship between the ERN and PES for efficient control of task performance. It was thus hypothesised that there would be a negative correlation between ERN amplitudes and PES, indicating that more negative error-related neural signals correspond to larger slowing of reaction times post-error (Hypothesis 3).

As relatively poorer motor skills in a healthy population may be associated with anxiety, it was hypothesised that motor proficiency scores would correlate with anxiety scores (Hypothesis 4). Based on previous suggestions linking control processes with anxiety, it was expected that both ERN and PES would correlate with anxiety (Hypothesis 5). Specifically, it was expected that ERN would correlate with anxiety in the negative direction suggesting higher anxiety in relation to more negative ERN amplitudes. PES would correlate with anxiety in the positive direction, suggesting that larger slowing of reaction times corresponds to higher anxiety levels.

## 2. Method

### 2.1 Participants

Participants were recruited by opportunity sampling either via the University of Surrey’s research volunteer system or by word of mouth. First and second year psychology undergraduate students were offered two research tokens for participation which are required within their degree. Additional participants were postgraduate students or individuals working at the university. All participants were entered into a prize draw of two £50 shopping vouchers. Exclusion criteria comprised individuals below the age of 18 years old and/or those who had a diagnosis of neurodevelopmental, psychiatric or neurological disorders. A total of 40 participants were recruited.

A median split method was used to allocate participants into two groups with higher or lower motor scores using the total raw scores on the Bruininks-Oseretsky Test of Motor Proficiency 2-short form (BOT2-SF; Bruininks & Bruininks, 2005). The scores ranged from 62 to 79 with the mean of 71.48 and the median of 72. Therefore, 23 participants with a score of 72 or more where placed in a high motor proficiency (HMP) group and 17 participants with a total score of 71 or less were placed in the lower motor proficiency group (LMP). In this group, six participants’ motor proficiency scores fell below the 10^th^ percentile indicating clinically significant motor difficulties and a possibility of undiagnosed DCD. The percentile calculations were based on age appropriate scale scores up to the age of 21. As no more scale scores are available beyond this age, the remaining calculations were based on the scale scores for 21-year olds. Some participants’ raw scores equated to the mean/median score and so the resulting two groups were uneven. It has been suggested that, in such situations, the power of the study may be reduced because many participants with similar scores are allocated to just one of the groups, which affects the obtained means and standard deviations on all variables (Iacobucci, Posavac, Kardes, Schneider, & Popovich, 2015). However, the study included group analyses as well as correlational analyses on all participants and thus it was important to include individuals representing the whole range of motor skills in order to test linear relationships between the investigated variables. The two resulting groups were significantly different on their motor proficiency, as indicated by an independent samples *t*-test (*t*(38)=8.06, *p*<.001, *d*=2.58). Demographic details for the overall sample as well as the separate groups are displayed in Table 1.

**Table 1.** Demographic information for participants in the overall sample and the two motor proficiency groups.

|  | Overall Sample | HMP | LMP |
| --- | --- | --- | --- |
| *N* | 40 | 23 | 17 |
| Age in years | 18-52 (M=26.2,<br>SD=9.0) | 18-52 (M=28.0,<br>SD= 9.7) | 18-49 (M=23.7,<br>SD=7.3) |
| Female Participants (%) | 60 | 52 | 76 |
| Left-handed (%) | 8 | 4 | 12 |
| Student (BSc, MSc, PhD; %) | 83 | 74 | 95 |
| **Ethnic Background** |  |  |  |
| White (%) | 82.5 | 87 | 76 |
| Black (%) | 7.5 | 4 | 12 |
| Asian (%) | 7.5 | 9 | 6 |
| Mixed (%) | 2.5 | 0 | 6 |

### 2.3 Materials

#### 2.3.1 Motor functioning

The Bruininks-Oseretsky Test of Motor Proficiency (2^nd^ Edition – short form; BOT2 -SF) was used to assess participants’ motor proficiency. BOT2-SF is a standardised test with normative scores provided for ages 4-21. There are currently no standardised measures to test motor proficiency in adults over 21. BOT2-SF was chosen because its validity and reliability in participants up to the age of 21 is stronger in comparison to other available measures (Hands, Licari, & Piek, 2015; McIntyre et al., 2017). It also has good test-retest reliability (Yoon, Scott, Hill, Levitt, & Lambert, 2006). In the study of motor functioning, it is often the case that standardised assessments designed for children and adolescents are used with adults where there is no equivalent age-appropriate measure. BOT2-SF and its components have previously been used in the study of adult populations (Du, Wilmut, & Barnett, 2015; Sahlander, Mattsson, & Bejerot, 2008; Silva et al., 2017). It consists of 12 items divided into four different motor domains: fine manual control, manual coordination, body coordination, and strength and agility. In the strength and agility domain, the researchers make a choice with regards to the type of push-up participants complete as part of the test. In this study, knee push-ups were chosen to accommodate for participants who were not used to engaging in strenuous exercise. The raw scores were totalled and used in the analysis because there were no standardised scores for participants over the age of 21. Higher scores indicate better motor proficiency. There is no agreement across the literature as to whether the raw scores should be used or whether the scale scores should be calculated based on the 21 years old category for all participants above that age. Both solutions are commonly used (Du et al., 2015; Sahlander et al., 2008; Silva et al., 2017).

#### 2.3.2 Cognitive measures

EEG recording was conducted during an arrow version of the Flanker task (Eriksen & Eriksen, 1974). The Flanker task was presented using E-Prime 2.0 software (Psychology Software Tools, 2012). Each trial consisted of seven arrowheads presented in Arial 24pt at the centre of the screen. The middle arrowhead was the target. Participants were instructed to press the letter “C” if the middle arrowhead was pointing left and the letter “M” if the arrowhead was pointing right on a standard computer keyboard. The three arrowheads positioned on each side of the target were distractors. The position of the distractor arrowheads was dependent on the test condition. In the congruent condition, the distractor arrowheads were pointing in the same direction as the target and in the opposite direction in the incongruent condition. A neutral condition was also included where distractor arrowheads were pointing down, resembling the letter “V”. Each trial was preceded by a fixation cross. The task was composed of 600 trials, including 200 trials in each condition, pseudorandomised across four blocks of 150 trials. Participants were instructed to take a break between each block and rest their eyes to avoid excessive blinking throughout the procedure. The time allowed to make a response in each trial was 600ms and between-trial intervals were jittered between 400 and 1600ms to prevent rhythmical responses. Figure 1 shows a diagram of the task conditions as presented to the participants.

**Figure 1.**
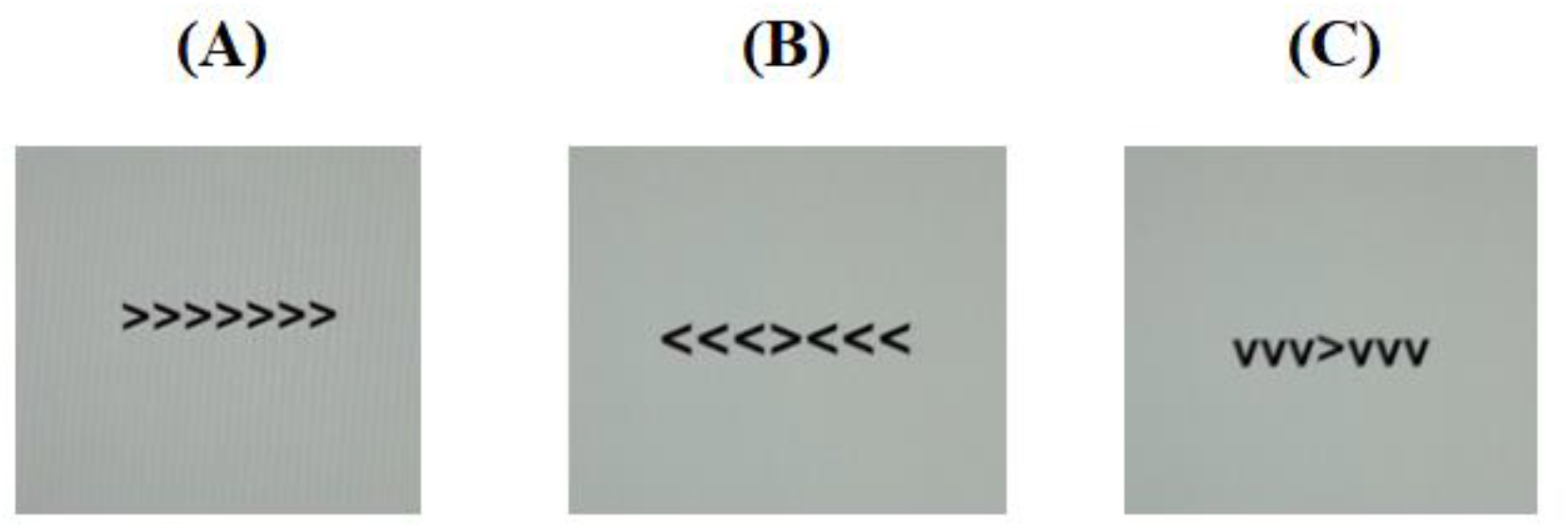
*(single column fitting)* An example of the conditions presented in the flanker task. (A) is the congruent condition, (B) is the incongruent conditions and (C) is the neutral condition.

Apart from task accuracy and reaction times, PES was also extracted as a behavioural measure using the robust method proposed by Dutilh et al. (2012). The method was chosen because it produces more reliable results in cases where reaction time variability may fluctuate throughout the task. It was expected that this could be the case in a Flanker task with three conditions of different difficulty (congruent, incongruent, neutral) and participants were expected to respond slower in the incongruent compared to the congruent condition. A correction was applied to the PES formula proposed by Dutilh et al. (2012) for this specific study where any omission errors were removed from the calculation because they differ from the commission errors in the pattern of response and may lead to post-omission speeding (Huang, Shum, & Chan, 2013). As a result, the PES was calculated in a pairwise approach around each commission error using the following formula:

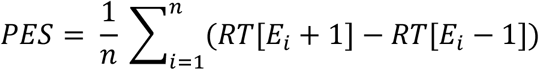

where *n* is the total number of commissioned errors, E_i_ is the i^th^ commission error trial, RT[E_i_-1] and RT[E_i_+1] are reaction times of the preceding and the subsequent correct trial, respectively. It was also required that E_i_-2 was a correct response because otherwise E_i_-1 (included in the formula) would have been a post-error RT and could bias the calculation.

#### 2.3.3 Neurophysiological measures

The aim of the EEG measurement was to obtain the error-related negativity (ERN) signal. EEG data were acquired using the BrainCap (EasyCap system kit, Neurospec) with 32 Ag/AgCl sintered electrodes in an extended 10/20 system. Impedance was kept below 5kΩ. Vertical and horizontal eye movements were recorded by electrodes located below and on the side of the left eye. Linked mastoid electrodes were used as the online reference and the ground electrode was located at the AFz location. Data were digitised at a sampling rate of at 500Hz. Offline analysis was carried out using BrainVision Analyser 2 (Brain Products, 2012). A bandpass filter 0.5-40 Hz was applied to the raw data. Eye movement correction was applied using the automatic independent component analysis approach for ocular correction following the default values across the whole dataset. Segments were sourced separately for correct and incorrect responses across all conditions within −150ms and 200ms of the response. Additional artefacts were rejected following the default values for the gradient, difference of values in intervals, minimal and maximal amplitude, and low activity. Baseline correction was applied at 150ms to 100ms prior to the response before the respective segments were averaged. Only participants with 8 or more artefact free error trials were included for ERN analyses, given previous suggestions that a minimum of 6-8 trials should be included for good internal consistency (Olvet & Hajcak, 2009). This criterion resulted in the exclusion of two participants in the HMP group from all analyses involving ERN. The error count in the retained sample ranged from 11 to 73 with the mean of 33.18 across the whole sample and no significant difference between the two groups (HMP *M*=31.19, LMP *M*= 35.65, *t*(36)=0.82, *p*=0.419, *d*=0.27). Only recordings from the Cz channel were used to measure the ERN. ERN mean amplitudes were extracted for a 50ms time window from 10 – 60ms post response separately for correct and incorrect responses. The respective trials were pooled together from the three conditions – congruent, incongruent and neutral – as most participants did not commit enough errors to allow for a comparison between conditions. All omission trials were excluded as omission-related brain activity has been reported to be different to commission-error related activity and primarily linked to attentional processes (Perri, Spinelli, & Di Russo, 2017; Yokota, Soshi, & Naruse, 2019). A difference wave was calculated to represent the ERN by subtracting the observed amplitude on the trials with correct responses from those with incorrect responses. This is an efficient method that has been shown to reduce the overlap with other ERPs (Luck, 2014) and is more robust in comparison to other available techniques, such as using the mean amplitude following incorrect responses only or the difference between the post-error negativity and the proceeding positivity (Fischer, Klein, & Ullsperger, 2017, Luck, 2014).

#### 2.3.4 Trait Anxiety

The Penn State Worry Questionnaire (PSWQ; Meyer, Miller, Metzger, & Borkovec, 1990) was used to measure the levels of trait worry to reflect participants’ trait anxiety tendencies. The scale has been validated to be a suitable tool for the investigation of the nature of worries as a trait (Olatunji, Schottenbauer, Rodriguez, Glass, & Arnkoff, 2007). It comprises 16 items which are answered on a 5-point Likert scale where 1 indicates “not at all typical of me” and 5 indicates “very typical of me”. A maximum score on the questionnaire is 80. Ascending scores reflect higher anxiety and a score of 60 or above reflects high levels of worry compared to normative samples (Gillis, Haaga, & Ford, 1995). The test has been shown to successfully distinguish participants with a generalised anxiety disorder (GAD) who experience excessive worry from other forms of anxiety such as social anxiety or obsessive compulsive disorder (OCD; Brown, Antony, & Barlow, 1992; Fresco, Mennin, Heimberg, & Turk, 2003). Thus, it can be used as a measure of anxiety reflecting traits of excessive worry (Fresco et al., 2003). This is relevant to the current study because participants with different sensitivity to error-making may be more or less worried about their performance (Frank et al., 2005; Holroyd & Umemoto, 2016). The PSWQ has previously been used in the study of the association between anxiety and error-related cognitive control in the general population (Hajcak, McDonald, & Simons, 2003; Pajkossy, Keresztes, & Racsmány, 2017).

### 2.4 Procedure

Ethical approval was obtained from the Faculty of Health and Medical Sciences Ethics Committee at the University of Surrey. Participants received the study information sheet when first expressing interest in the study and then again before signing the consent form when arriving in the room set up for the procedure. All participants were aware of the nature of the study. Namely, they were told that the relationship between their motor skills, anxiety/worry and cognitive control processes would be studied. After signing the consent form, they were asked to complete the demographic and PSWQ questionnaires. Then, they were taken to an adjacent room where they completed the BOT2-SF tasks. Finally, they were taken to a smaller EEG room where the EEG cap was applied. For each participant, the scalp was cleaned where the electrodes were placed, and an electrolyte paste was used for conductance. Once the setup was finished, participants completed the Flanker task. On average, the duration of the procedure was 1 hour and 45 minutes including breaks. The whole procedure was conducted in the same order by the same researcher for each participant. Participants were debriefed after the study to further clarify that the EEG analysis would focus on the instances when they made errors during the task. This time was also used to answer any questions.

### 2.5 Data Analysis

Statistical analysis of the data was conducted using SPSS 24 (IBM, 2016). All variables used to test the hypotheses were tested for normality in their distribution. Age and PES were log-transformed due to significant skew and kurtosis. A screening process included a correlational investigation of the likely effects of sample characteristics (age and gender) on the dependent variables including ERN, PES and anxiety and motor functioning. There was a significant positive correlation between age and motor proficiency (*r*=.435, *p*=.005).

For the analysis of the flanker effects, the whole sample of 40 participants was included (HMP *N*=23, LMP *N*=17). The behavioural results included reaction times (RT), with omission trials excluded, and accuracy (proportion correct responses), where omission trials were retained. This was collected for both groups across the three conditions. The differences between flanker conditions (congruent, incongruent and neutral) and motor proficiency groups (HMP and LMP) were tested using a two-way mixed ANOVA.

For the investigation of cognitive control processes on the neural level, the ERP difference wave mean amplitudes were compared between the two motor proficiency groups at the Cz channel using a two-tailed independent samples *t*-test. Two participants from the HMP group were excluded from this analysis as they committed fewer than eight errors during the task, resulting in the following group sizes, HMP *N*=21, LMP *N*=17. These two cases were also excluded from all following analyses with ERN as a variable. All cases were checked for outliers using z-scores and none were identified.

For the investigation of behavioural adaptation, a two-tailed independent samples *t*-test was used to test the difference between the mean PES scores for both groups. In this analysis, and all further analyses of the PES, a total of 11 participants were excluded, resulting in the following group sizes, HMP *N*=17, LMP *N*=12. Ten of these cases were removed because participants committed fewer than 11 errors which were suitable for PES analysis, as calculated with the formula presented above. This resulted in 30 remaining cases with the range of qualifying errors between 11 and 55 and a mean of 25.63. The mean of qualifying PES errors did not significantly differ between the two groups when tested with an independent samples *t*-test (HMP *N* 24.18, LMP *M*= 27.54, *t*=0.24, *p*=.467, *d*=0.27). Another case was removed which was an extreme outlier with a large number of committed errors (*z*=2.39). It was identified after the initial exclusion of the ten cases with too few errors. The specificity of error trials suitable for the analysis of post-error behaviours makes it challenging to obtain a high number of erroneous trials, which could introduce bias when testing hypotheses due to low power. Danielmeier and Ullsperger (2011) suggest that experimenters should exercise caution, but so far there is no recommendation for the minimal number of erroneous trials that should be included in PES analyses. In comparison to ERN, reaction time data tends to be more variable and it is suggested that as many trials as possible should be included to obtain a representative mean for each participant (Danielmeier & Ullsperger, 2011). In the current experiment, it was identified that 25% of participants committed fewer than 11 errors qualifying for the analysis of PES. It was not feasible to raise the threshold of 11 trials any higher as this would lead to many more exclusions and the current analysis would be underpowered and not sensitive enough to identify the effects of interest. Before commencing group analyses, all included cases were checked for outliers using z-scores and none were identified. Lastly, the relationship between ERN and PES was measured using a Pearson correlation.

For the correlational analyses involving anxiety, a one-tailed partial correlation, controlling for age, was conducted to test the relationship between motor skills and anxiety. In this analysis, one multivariate outlier was removed, and the total of included cases was *N*=39. Age was included as a controlled variable because a significant relationship with motor skills had been identified during the screening of the variables.

Subsequently, one-tailed partial correlations, controlling for motor skills, were conducted to test the relationship between anxiety and ERN as well as anxiety and PES. Motor proficiency was partialled out due to the expected relationship between motor skills and anxiety

#### 2.5.4 Exploratory data analysis

Further analyses were informed by the outcomes from the hypothesis-driven analyses. These focused on participants’ accuracy and the type of errors they made to investigate whether cognitive control and behavioural adaptation helped participants to perform accurately and avoid errors. Correlational analyses were used to test the relationship between ERN and accuracy and PES and accuracy. Additionally, participant errors were divided into omissions and commissions and the differences between these error rates where compared for both motor proficiency groups using an independent *t*-test analysis. This was decided because omission errors could be associated with PES due to the slowing of reaction times and a consequent inability to execute the responses in time. All analyses were Bonferroni corrected.

## 3. Results

### 3.1 Flanker Effect

As predicted in Hypothesis 1, both groups displayed typical patterns of responses for a flanker effect, including reduced accuracy and longer reaction times in the incongruent trials compared to congruent and neutral trials. This reflects the commonly observed cognitive conflict during the incongruent trials. Accuracy and reaction times across conditions are displayed in Table 2.

**Table 2.**
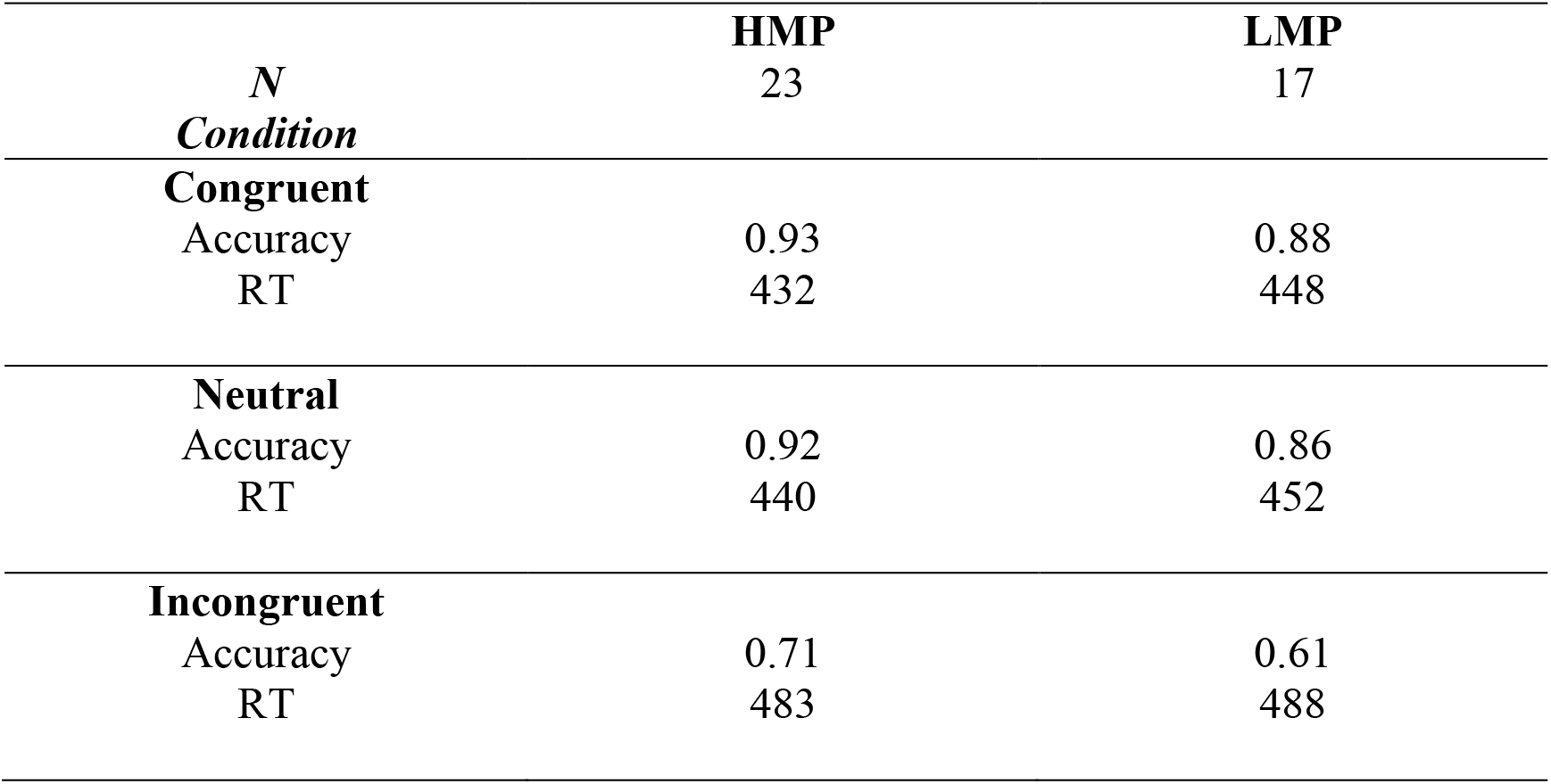
Accuracy proportion and reaction times (ms) on the flanker task for each motor proficiency group.

For accuracy, a two-way (flanker condition, motor proficiency group) mixed ANOVA with applied Greenhouse-Geisser correction revealed a significant effect of condition, *F*(1.14, 76)=100.40, *p*<.001, *η_p_^2^* = .73. The effect of motor proficiency group was also significant, *F*(1, 38)=6.77, *p*=.013, *η_p_^2^* = .15, with the LMP group scoring below the HMP group overall. There was no significant interaction between group and condition, *F*(1.14, 76) =1.31, *p*=.264,, *η_p_^2^* = .03. This indicates that there was a significant general effect of the flanker condition on accuracy in the expected direction which did not differ between the two groups. However, task accuracy in general significantly differed between the two motor proficiency groups with higher accuracy rates in the HMP group.

For reaction times, a two-way (flanker condition, motor proficiency group) mixed ANOVA with applied Greenhouse-Geisser correction also revealed a significant effect of condition for RTs, *F*(1.21, 76)=143.9, *p*<.001, *η_p_^2^* =.79. However, the group effect was not significant, *F*(1, 38)=1.44, *p*=.237, *η_p_^2^* = .037 and neither was the interaction *F*(1.21, 76)=1,628, *p*=.203, *η_p_^2^* = .041. The differences between the RTs obtained for each condition were assessed for the whole sample using post-hoc t-tests which confirmed that RTs were longer for incongruent compared to congruent and neutral trials. This indicates that there was a significant general effect of the flanker condition on RTs which did not differ between the two groups.

### 3.2 Cognitive control and behavioural adaptation

To address Hypothesis 2, the difference in the ERN between HMP (*N*=21, *M* = −11.45, *SD*=7.63) and LMP (*N*=17, *M* = −6.76, *SD*=5.24) groups at the Cz channel was tested using a two-tailed *t*-test with Welch correction. In contrast to the predictions, the HMP group had significantly higher ERN *t(35)*=2.24, *p*=.031, *d*=0.71 than the LMP group. Figure 2 shows the obtained post-error, post-correct and difference waves at the Cz as well as the topographic distributions of the difference waves for both groups.

**Figure 2.**
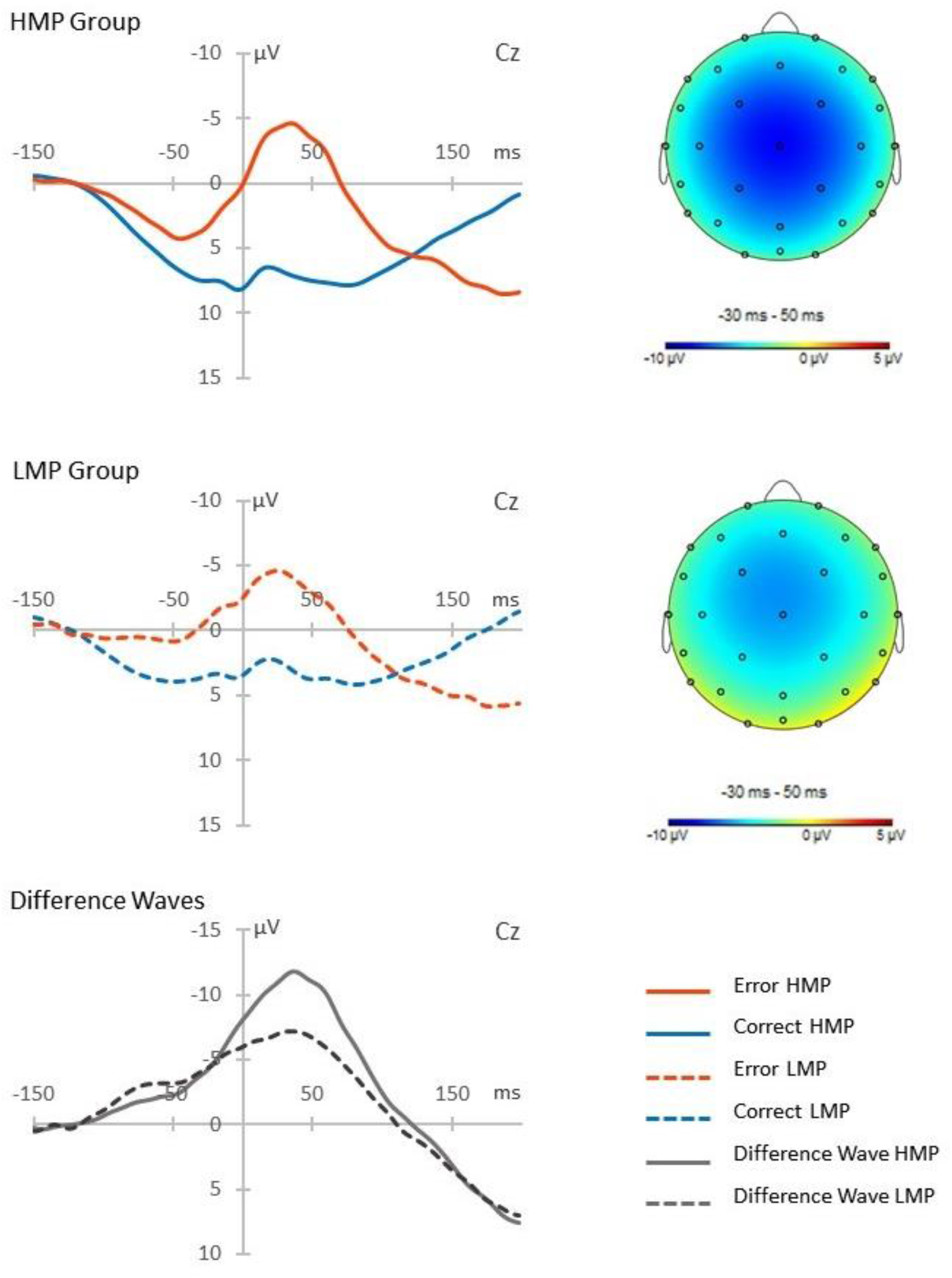
*(double column fitting)* Post-error and post-correct waveforms presented for each motor proficiency group. The difference waves reflect the ERN response which is additionally presented with a topographic distribution.

Additionally, a two-tailed *t*-test with Welch correction was conducted on the log-transformed PES values between the HMP (*N*=17, *M*=1.30 *SD*=0.36) and LMP (*N*=12, *M*=1.56, *SD*=0.21) groups. The LMP group had significantly longer PES than the HMP group, as expected, *t*(26.19)=1.40, *p*=.021, *d*=0.88. Figure 3 shows a raw descriptive-inferential graph for the group differences on ERN and PES.

**Figure 3.**
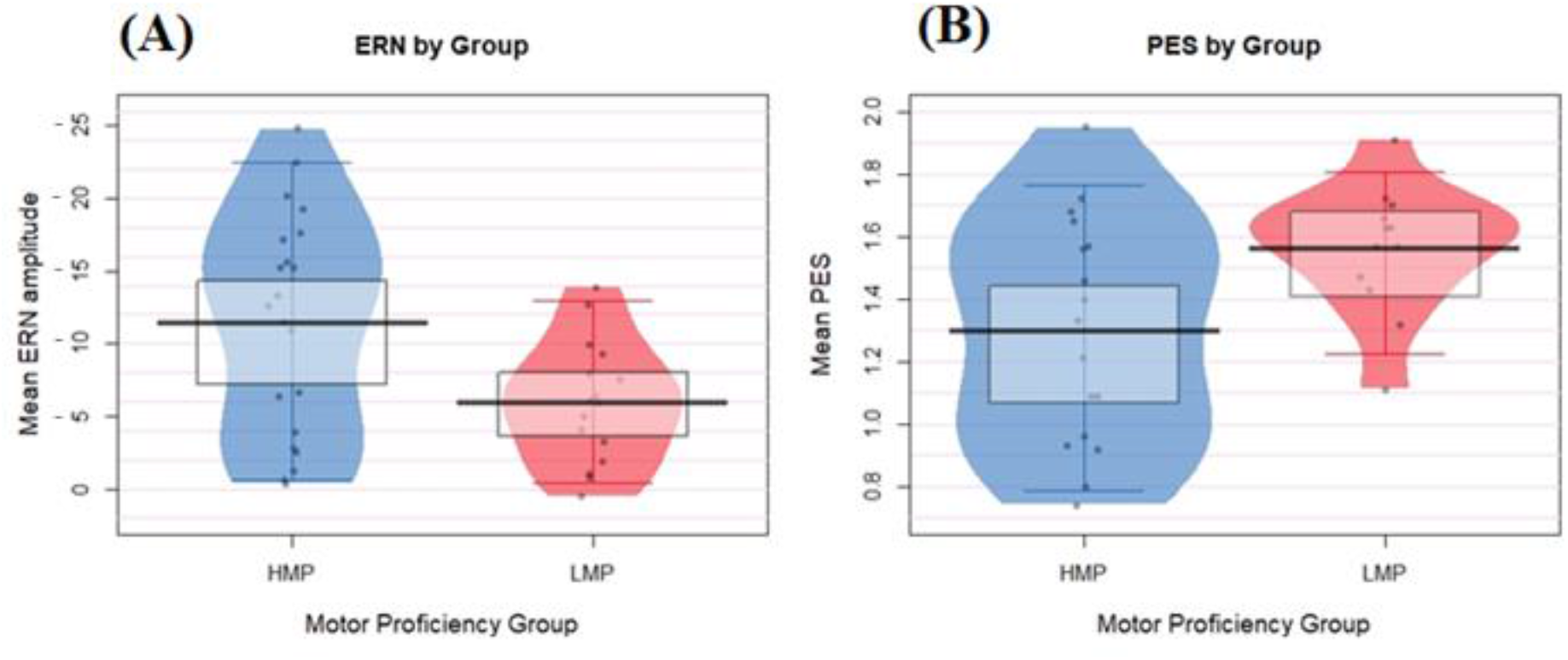
*(double column fitting)* Group data is displayed in the form of jittered dots and data density, indicated by the width of the shape. The mean value is marked by the black indicator line and the box around it marks the high-density intervals of the mean. (A) Differences in the ERN values between the two groups. ERN values were inverted from negative to positive values to represent larger ERN at the top of the plot and smaller ERN at the bottom. (B) Differences in the PES values between the two groups.

Contrary to Hypothesis 3, the correlation between the ERN and PES was not significant, *N*=29, *r*=−0.041, *p*=.833.

### 3.3 Cognitive control, behavioural adaptation and anxiety

In line with Hypothesis 4, the partial correlation between anxiety and motor proficiency, controlled for participant age, was significant, *r*=−.360, *p*=.027. This suggests that anxiety is associated with participants’ motor proficiency. Figure 4 presents the scatter plot for this relationship. However, in contrast to Hypothesis 5, the partial correlation between ERN and trait anxiety, controlled for motor skills, was not significant, *N*=38, *r*=.19, *p*=.13. Similarly, the partial correlation between PES and trait anxiety was also not significant, *N*=29, *r*=.251, *p*=.198.

**Figure 4.**
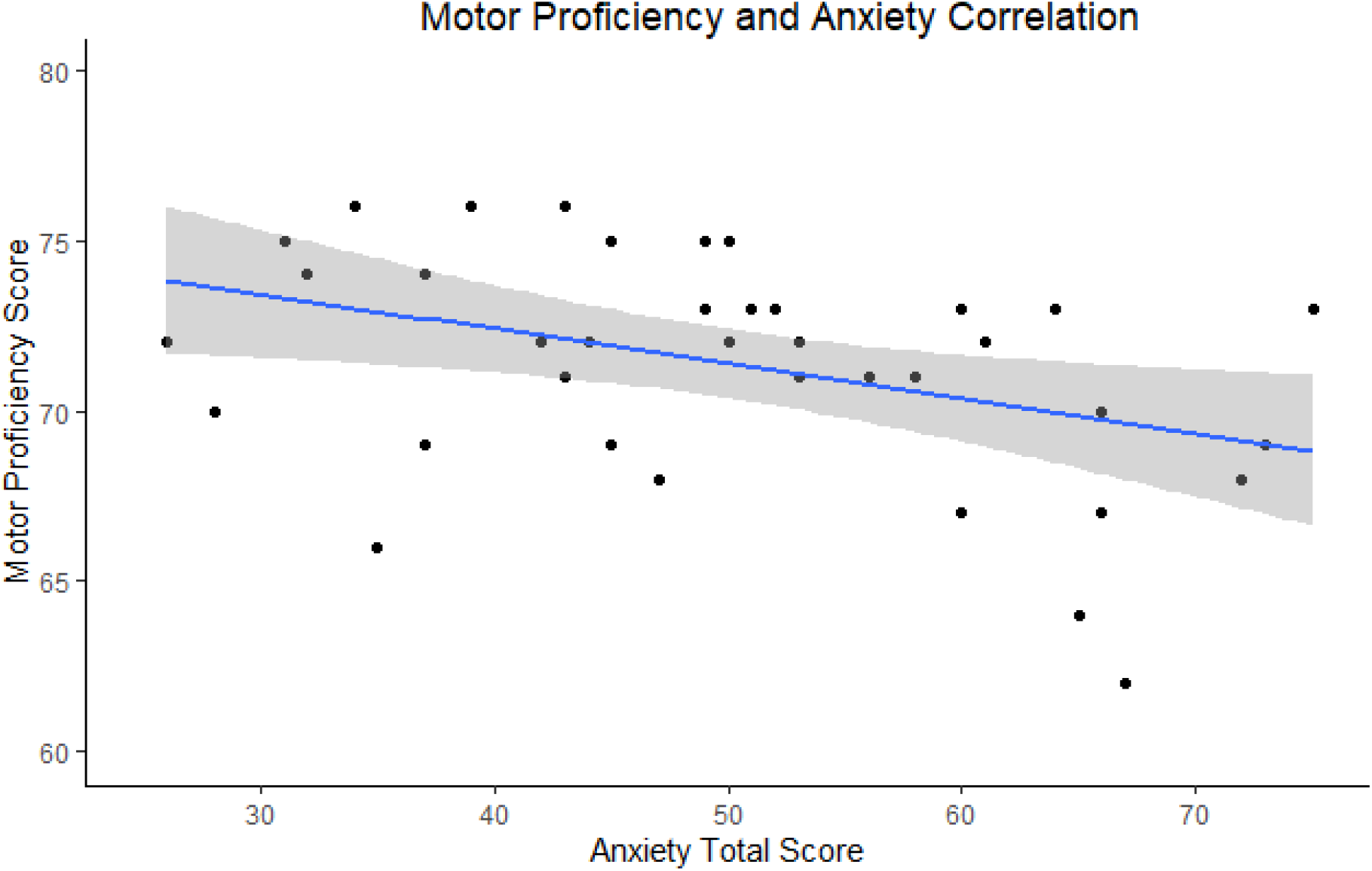
*(single column fitting)* Visual representation of the relationship between anxiety scores (x axis) and motor proficiency (y axis) with a marked line of best fit and the confidence intervals reflected with the shadowed area.

### 3.4 Exploratory Analyses

Accuracy and error rate variables were tested in relation to ERN and PES to investigate the efficiency of cognitive control and behavioural adaptation in individuals with different levels of motor proficiency. Accuracy significantly correlated with ERN in the negative direction (*N*=38, *r*=−.398, *p*=.026), indicating that higher accuracy is associated with larger ERN. There was no significant correlation between accuracy and PES (*N*=29, *r*=−.051, *p*=.791). The results therefore suggest that cognitive control processes on the neural level may facilitate participants’ performance, but this may not be the case for behavioural adaptation. Figure 5 presents the scatterplot for the relationship between ERN and accuracy. The omission and commission rates were extracted for both motor proficiency groups; the descriptive statistics are provided in Table 3. They were compared between the two groups using independent *t*-tests, revealing that the LMP group made significantly more omissions compared to the HMP group (*t*(38)=2.40, *p*=.044, *d*=0.77). There was no difference for commission rates (*t*(38)= 1.21, *p*=.466, *d*=0.39).

**Figure 5.**
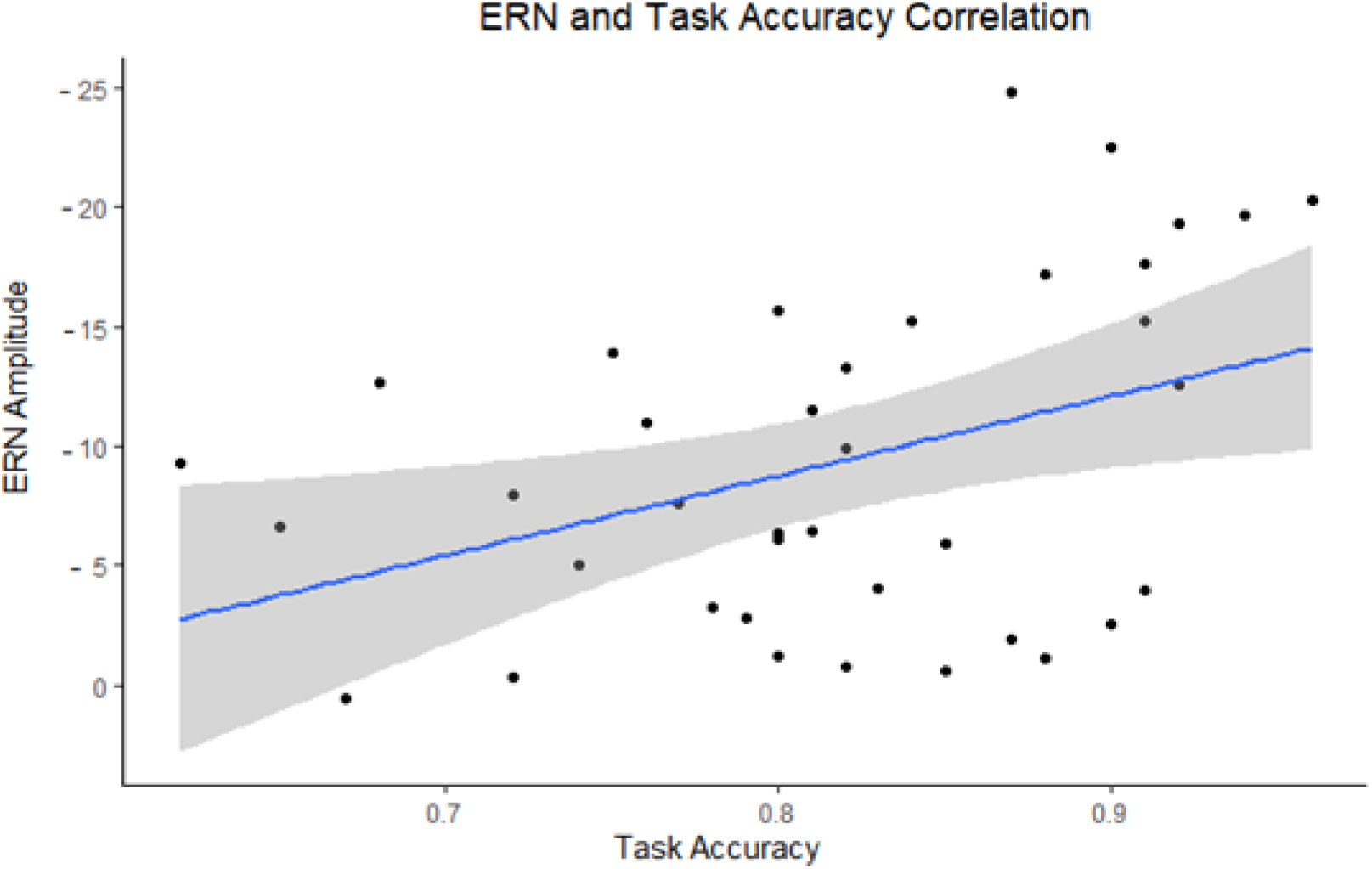
*(single column fitting)* Visual representation of the relationship between task accuracy (x axis) and ERN amplitudes (y axis) with a marked line of best fit and the confidence intervals reflected with the shadowed area. The ERN values have been reversed for visualisation purposes as the more negative the ERN the stronger the cognitive control signal.

**Table 3.** Means and standard deviations for omission and commission rates for the two motor proficiency groups.

| Group | Omission Errors | Commission Errors |
| --- | --- | --- |
| HMP | 59.74 (40.8) | 29.00 (16.4) |
| LMP | 93.65 (48.7) | 35.65 (18.2) |

## 4. Discussion

The current study aimed to investigate the motor-cognitive interaction in the general population by focusing on cognitive control and behavioural adaptation in individuals with different levels of motor proficiency. It also investigated whether these mechanisms were associated with anxiety.

### 4.1 Task performance, cognitive control and behavioural adaptation

First, in order to assess the cognitive control and behavioural adaptation mechanisms, healthy adult participants were split into two groups based on their motor ability including high motor proficiency (HMP) and low motor proficiency (LMP). They completed the flanker task during an EEG recording. Both groups demonstrated longer reaction times and lower accuracy in the incongruent condition of the task. Such a pattern of performance is expected in the flanker task. It has been regularly reported in previous studies and indicates that the task is completed correctly, which is a prerequisite for conducting further analyses (Fischer, Danielmeier, Villringer, Klein, & Ullsperger, 2016; Grützmann, Endrass, Klawohn, & Kathmann, 2014). However, the observed group effect on task-accuracy, wherein the LMP group performed worse than the HMP group, was not expected. This finding is reasonable given that individuals with lower motor proficiency may not be able to complete a task that requires motor responses to the same standard as individuals with higher motor proficiency. A third of the participants in the LMP group had motor difficulties of clinical significance, which may suggest cases of undiagnosed DCD within the group. In this case, it is unsurprising that they found the task more challenging and performed less accurately.

However, previous research found that participants with motor disorders including DCD and Tourette’s syndrome completed reaction-time-based cognitive tasks with accuracy that was not different to that of the control group (Querne et al., 2008; Schüller et al., 2018). This led to the assumption that cognitive control and behavioural adaptation strategies must help individuals with motor disorders to compensate for their motor difficulties and perform the tasks as well as healthy individuals. Therefore, a similar outcome was expected in the current study. It should be noted that although both Querne et al. (2008) and Schüller et al. (2018) observed no statistical significance in the difference between the accuracy of the clinical and control groups, without equivalence testing or Bayesian analysis this is not evidence for equivalent task performance. Therefore, it does not preclude that there might be differences in other cohorts. The current study suggests that task accuracy may differ between individuals with better and worse motor ability, although this evidence is provided in the context of individuals without diagnoses of motor disorders. The issue of task accuracy is an important point in this type of research as it directs the interpretation of the findings reflecting cognitive control and behavioural adaptation mechanisms.

In the current study, cognitive control was indexed by the ERN signal, while behavioural adaptation was indexed by the reaction-time based PES. It was found that the LMP group achieved lower task accuracy and had significantly lower mean ERN signal, but their PES was larger compared with the HMP group. The ERN finding suggests that individuals with poorer motor proficiency may have attenuated activity in the ACC or more varied latency of the error-related neural response in comparison to individuals with better motor proficiency. This contradicts the claims by Querne et al. (2008) and Schüller et al. (2018), who suggest that ACC activity may be enhanced in individuals with motor difficulties in order to facilitate task accuracy as a compensation for their motor difficulties. One explanation for the contradicting results could be that individuals with a diagnosis of a neurodevelopmental motor disorder such as DCD or TS are more likely to develop compensatory mechanism to facilitate their performance despite experienced difficulties. Both DCD and TS are neurodevelopmental disorders affecting individuals since childhood, thus giving an opportunity to develop compensatory strategies over many years and in a range of different situations and activities. They may receive therapy or input to help them manage their difficulties, thus there is generally more support for their development of compensatory processes which may consolidate on the neural level. In comparison, individuals who have no motor disorders but present with poorer motor functioning may not see their motor performance to be problematic and they may not receive specialist input to help with the development of management strategies or compensatory mechanisms. Thus, they may not feel the need to compensate or monitor their performance with any additional effort. It is also possible that the M-C interaction may be qualitatively different in individuals with motor disorders compared to healthy individuals with relatively poorer motor functioning. In fact, the M-C interaction may be syndrome-specific and operate differently for different types of motor disorders, which may explain why results from the general population do not reflect those from studies on motor disorders.

Our initial hypothesis that the PES would be larger in the LMP group than the HMP group was supported by the analysis, albeit in concurrence with an unexpected attenuation of the ERN. High PES and attenuated ERN may again suggest that individuals with lower motor proficiency cannot successfully employ cognitive control mechanisms to facilitate their performance as they are not aware of the need for it. However, because their motor skills are below average, they may still struggle with the completion of certain tasks where they adapt their performance behaviourally in order gain better control over their accuracy. This technique may not be so effective because it is based solely on the slowing of reaction times to avoid further errors. There is a lack of cognitive control engagement which would help to monitor and update the pattern of responses for the duration of the task, which would be a more efficient strategy. These findings suggest that the ACC compensatory processes developed by individuals with neurodevelopmental motor disorders do not reflect a general mechanism of the M-C interaction. Such compensation is not observed in individuals with a sub-clinical level of motor difficulties. It may potentially develop over many years, across a range of situations and may require that an individual is aware of their difficulties and engages conscious effort into trying to improve their performance.

In summary, analyses of the flanker task revealed that individuals with lower motor proficiency had lower task accuracy, had more attenuated cognitive control during the task but also used more effort in the behavioural adaptation of their responses. On the other hand, individuals with higher motor proficiency had higher accuracy, relatively enhanced cognitive control and did not rely on adapting their performance behaviourally to the same extent as the LMP group. Contrary to previous research on ERN and PES, the current study found no evidence for a relationship between these two compensatory mechanisms. Questions remain with regards to whether the investigated cognitive control and behavioural adaptation mechanisms help to facilitate task performance. Therefore, in an exploratory analysis, the correlation between task accuracy was investigated in relation to the ERN and PES to understand whether these mechanisms facilitate task performance and could explain the accuracy difference between the two groups.

As a result of the exploratory analyses, we found a significant medium-large correlation between task accuracy and ERN, but the correlation between PES and task accuracy was not significant. This suggests that ACC cognitive control processes, as reflected by the ERN, facilitate successful task performance. This effect has been shown in previous research (Themanson, Rosen, Pontifex, Hillman, & McAuley, 2012) and is consistent with the models of cognitive control and the proposed patterns of interaction between motor and cognitive networks within the ACC (Brown & Alexander, 2017; Holroyd & Coles, 2002). It is therefore possible that the HMP group benefitted from strong cognitive control processes and thus achieved better task accuracy compared to the LMP group. On the other hand, the study did not find evidence that behavioural adaptation in the form of PES may contribute to the improvement of task accuracy, given that the tested correlation produced a very negligible effect size of *r*=−.051, which was not statistically significant. It may be the case that the extended reaction times occurring after error commission have led participants in this group to make omission errors. They slowed their reaction times to the extent where they were not able to execute their prepared response in time before the next trial began. To explore this possibility, the rate of commission and omission errors was compared between the two groups and it was found that the LMP group made significantly more omissions but not commissions compared to the HMP group. This suggests that their application of behavioural adaptation in the form of slowing of reaction times might have been counterproductive and could have contributed to the difference in task accuracy between the two groups. This reflects that participants in the LMP group prioritised accuracy over speed, perhaps considering omission errors as less indicative of their ability to perform the task correctly. It may be the case that, if they were given a longer time window in which to make their responses, their accuracy would have been comparable or even better than that of the HMP group. This would be the result of their application of the PES as a strategy which is known to be a self-regulatory behavioural control mechanism characteristic of individuals who obtain high task accuracy (Steinborn, Flehmig, Bratzke, & Schröter, 2012). It would be interesting to assess to what extent PES is employed by individuals with DCD and TS and whether it may complement enhanced ERN as an optimal engagement of cognitive and behavioural control in the context of the M-C interaction.

### 4.2 Anxiety, cognitive control and behavioural adaptation

Cognitive control (ERN) and behavioural adaptation (PES) were both tested for their relationship with anxiety following previous reports linking enhanced control mechanisms with increased anxiety levels (Hajcak et al., 2003). This investigation is important due to the predisposition of individuals with poorer motor functioning to experiencing increased anxiety (Rigoli et al., 2017). The current study found a significant relationship between motor proficiency and anxiety in line with previous research (Rigoli et al., 2017) thus providing a good opportunity to investigate whether this could be associated with participants’ sensitivity to error making and the need to apply cognitive control processes and behavioural adaptation mechanisms. The current study found no significant correlations between ERN or PES with anxiety. For ERN, this may be caused by the fact that it was attenuated in the LMP group, contrary to expectations. Rigoli et al. (2017) suggested an explanation for the increased anxiety observed in individuals with lower motor proficiency, rather than increased sensitivity to error-making, based on the Environmental Stress Hypothesis. They claim that individuals’ experiences of motor difficulties and difficult situations they may have to deal with which to anxious tendencies. It is possible that, in the current study, participants in the LMP group may have found the flanker task to be more stressful than the HMP group because the requirement to make fast motor responses was challenging. This task-specific stress may be related to task-specific control mechanisms such as the ERN or PES. However, anxiety levels were tested prior to the completion of the task, suggesting that anxiety in the current study was indicative of a trait, rather than a stress response to a task. If this were the case, then the lack of correlation between trait anxiety and task-specific ERN or PES is perhaps not surprising.

### 4.3 Implications

The current study suggests that individuals with lower motor proficiency may be less efficient in the application of cognitive control mechanisms to facilitate their task performance. Instead, they apply behavioural adaptation strategies which may be counterproductive, as demonstrated in the current study where the LMP group showed larger post-error slowing and an increased rate of omission errors. Therefore, individuals with poorer motor skills may appear to be less accurate or less successful when tested under fast-paced conditions. They may have a tendency to take a slower approach to tasks in order to ensure that they complete them correctly despite certain motor difficulties they experience, thus emphasising accuracy over speed. This may have important implications for educational and occupational settings for healthy individuals as well as individuals with health conditions affecting their motor control. Further research should aim to investigate cognitive control and behavioural adaptation mechanisms on tasks which resemble everyday activities to learn about the impact of fast-paced conditions in a more ecologically valid setting. For instance, in educational settings, children and young people with a range of motor skills must complete tasks under time pressure within the classroom. The current study suggests that those with poorer motor skills are likely to struggle under these conditions but could potentially perform well if given additional time.

In addition, the significant relationship between motor skills and anxiety should be considered. This suggests that even individuals with no motor disorders are at risk of developing anxious tendencies in relation to their motor functioning. This should be considered by health professionals and in the context of educational and occupational settings. Further research may aim to investigate how these risks may be mitigated or alleviated by certain adaptations such as the proposed extended time for task completion or more direct support to make tasks requiring motor activity less strenuous, thus requiring less adaptation.

It would also be of interest to investigate whether the ERN would still be attenuated or more varied in latency in individuals with lower motor proficiency when tested under slow-paced conditions compared to a fast-paced condition. Perhaps the engagement of enhanced error control mechanisms in this group is dependant on task difficulty. It could be possible to investigate the optimal conditions for the efficient engagement of cognitive control processes and successful task performance for individuals with lower motor proficiency in healthy and clinical populations, which may in turn inform practical solutions to support individuals with poorer motor functioning in everyday activities.

### 4.4 Limitations

The main limitation of the current study is the use of the median split method to create the two groups with different motor proficiency levels. Participants who were recruited reflected a wide range of motor skills with many cases nesting around the median score of 72, which made the two resulting groups uneven in size. This could have affected the power of the study and made it less likely to detect significant differences. If, instead, the two groups were oversampled to represent healthy individuals with high and low motor proficiency scores only, the differences between the groups would have been clearer and easier to detect by statistical analyses. However, as already mentioned, the study tested linear relationships between the observed variables and therefore it was important to recruit participants whose motor proficiency scores would form a continuous variable. In addition, the groups were significantly different on their motor proficiency level, with a large effect size of more than 2 standard deviations (*d*=2.58). In Figure 3, it is evident that the median split worked well to separate a homogenous group of participants with lower motor proficiency (LMP), which may be seen as a sub-clinical group with consistently small variance across their ERN and PES. In comparison, the HMP group was more heterogenous, with individuals often obtaining ERN or PES comparable to that of participants in the LMP group, which could be for reasons other than their motor proficiency. In general, the current study obtained a sufficient sample to detect the effects of interest despite the uneven split.

One additional disadvantage of the study is the possibility of sampling bias. A significant positive correlation between age and motor proficiency was found, suggesting that the older the participants the more likely they were to present with better motor skills. It is widely understood that motor skills decline as a function of age in the general population (Leversen, Haga, & Sigmundsson, 2012). It is possible that older individuals with relatively better motor skills than their peers were more likely to volunteer and take part in the study as they felt more confident about their performance. In the current study, all ANOVA and correlational analyses on motor proficiency were controlled for the effect of age. However, future research may aim to plan their recruitment strategy to be equally inviting for those who have different levels of motor proficiency across ages to avoid sampling biases.

Lastly, there are comparisons drawn between the results of this study and other studies conducted with participants with DCD and TS (Querne et al., 2008; Schüller et al., 2018). The comparisons are only relative considering that the methodology used in the current study was different. To support the conclusions of this study and the understanding of the M-C interaction in the context of healthy individuals as well as those with motor disorders, future research and any replications of the current project should aim to make direct comparisons between clinical and sub-clinical groups using the same methodology.

### 4.5 Conclusions

The lower motor proficiency group engaged cognitive control processes less efficiently than the individuals in the high motor proficiency group. Instead, they adapted their task performance behaviourally through slowing their reaction times, which was less effective. This suggests that the interaction between motor and cognitive processes in healthy individuals may not be comparable to that of individuals with motor disorders. The results of this study provide an important baseline for the understanding of the changes in motor-cognitive processes in clinical, sub-clinical and healthy populations.

## CRediT Authorship Statement

**Marta Topor:** conceptualisation, data curation, formal analysis, investigation, methodology, project administration, validation, visualisation, writing – original draft. **Bertram Opitz:** methodology, formal analysis, software, supervision, validation, visualisation, writing – review & editing. **Hayley Leonard:** conceptualisation, supervision, writing – review & editing.

## References

Bloch, M. H., & Leckman, J. F. (2009). Clinical course of Tourette syndrome. Journal of Psychosomatic Research, 67(6), 497–501. doi:10.1016/j.jpsychores.2009.09.002

Brown, T. A., Antony, M. M., & Barlow, D. H. (1992). Psychometric properties of the Penn State Worry Questionnaire in a clinical anxiety disorders sample. Behaviour Research and Therapy, 30(1), 33–37. doi:10.1016/0005-7967(92)90093-v

Brown, J. W., & Alexander, W. H. (2017). Foraging value, risk avoidance, and multiple control signals: How the anterior cingulate cortex controls value-based decision-making. Journal of Cognitive Neuroscience, 29(10), 1656–1673

Bruininks, R. H., & Bruininks, B. D. (2005). Bruinkinks-Oseretsky Test of Motor Proficiency, 2nd ed. Circle Pines, MN: NFER-Nelson.

Cairney, J., Veldhuizen, S., & Szatmari, P. (2010). Motor coordination and emotional–behavioral problems in children. Current opinion in psychiatry, 23(4), 324–329.

Chang, A., Chen, C.-C., Li, H.-H., & Li, C.-S. R. (2014). Event-related potentials for post-error and post-conflict slowing. Plos One, 9(6), e99909. doi:10.1371/journal.pone.0099909

Damiano, D. L., Zampieri, C., Ge, J., Acevedo, A., & Dsurney, J. (2016). Effects of a rapid-resisted elliptical training program on motor, cognitive and neurobehavioral functioning in adults with chronic traumatic brain injury. Experimental Brain Research, 234(8), 2245–2252. doi:10.1007/s00221-016-4630-8

Danielmeier, C., & Ullsperger, M. (2011). Post-error adjustments. Frontiers in psychology, 2, 233. doi:10.3389/fpsyg.2011.00233

Du, W., Wilmut, K., & Barnett, A. L. (2015). Level walking in adults with and without Developmental Coordination Disorder: An analysis of movement variability. Human Movement Science, 43, 9–14. doi:10.1016/j.humov.2015.06.010

Dutilh, G., van Ravenzwaaij, D., Nieuwenhuis, S., van der Maas, H. L. J., Forstmann, B. U., & Wagenmakers, E.-J. (2012). How to measure post-error slowing: A confound and a simple solution. Journal of mathematical psychology, 56(3), 208–216. doi:10.1016/j.jmp.2012.04.001

Eriksen, B. A., & Eriksen, C. W. (1974). Effects of noise letters upon the identification of a target letter in a nonsearch task. Perception & Psychophysics, 16(1), 143–149. doi:10.3758/BF03203267

Fischer, A. G., Danielmeier, C., Villringer, A., Klein, T. A., & Ullsperger, M. (2016). Gender Influences on Brain Responses to Errors and Post-Error Adjustments. Scientific Reports, 6, 24435. doi:10.1038/srep24435

Fischer, A. G., Klein, T. A., & Ullsperger, M. (2017). Comparing the error-related negativity across groups: The impact of error-and trial-number differences. Psychophysiology, 54(7), 998–1009. doi:10.1111/psyp.12863

Frank, M. J., Woroch, B. S., & Curran, T. (2005). Error-related negativity predicts reinforcement learning and conflict biases. Neuron, 47(4), 495–501. doi:10.1016/j.neuron.2005.06.020

Fresco, D. M., Mennin, D. S., Heimberg, R. G., & Turk, C. L. (2003). Using the Penn State Worry Questionnaire to identify individuals with generalized anxiety disorder: a receiver operating characteristic analysis. Journal of Behavior Therapy and Experimental Psychiatry, 34(3-4), 283–291. doi:10.1016/j.jbtep.2003.09.001

Fu, Z., Wu, D.-A. J., Ross, I., Chung, J. M., Mamelak, A. N., Adolphs, R., & Rutishauser, U. (2019). Single-Neuron Correlates of Error Monitoring and Post-Error Adjustments in Human Medial Frontal Cortex. Neuron, 101(1), 165–177.e5. doi:10.1016/j.neuron.2018.11.016

Gillis, M. M., Haaga, D. A., & Ford, G. T. (1995). Normative values for the beck anxiety inventory, fear questionnaire, Penn state worry questionnaire, and social phobia and anxiety inventory. Psychological Assessment, 7(4), 450.

Grützmann, R., Endrass, T., Klawohn, J., & Kathmann, N. (2014). Response accuracy rating modulates ERN and Pe amplitudes. Biological Psychology, 96, 1–7. doi:10.1016/j.biopsycho.2013.10.007

Hajcak, G., McDonald, N., & Simons, R. F. (2003). Anxiety and error-related brain activity. Biological Psychology, 64(1-2), 77–90. doi:10.1016/S0301-0511(03)00103-0

Hands, B., Licari, M., & Piek, J. (2015). A review of five tests to identify motor coordination difficulties in young adults. Research in Developmental Disabilities, 41-42, 40–51. doi:10.1016/j.ridd.2015.05.009

Hauser, T. U., Iannaccone, R., Stämpfli, P., Drechsler, R., Brandeis, D., Walitza, S., & Brem, S. (2014). The feedback-related negativity (FRN) revisited: new insights into the localization, meaning and network organization. Neuroimage, 84, 159–168. doi:10.1016/j.neuroimage.2013.08.028

Hill, E. L., & Brown, D. (2013). Mood impairments in adults previously diagnosed with developmental coordination disorder. Journal of mental health (Abingdon, England), 22(4), 334–340. doi:10.3109/09638237.2012.745187

Holroyd, C. B., & Coles, M. G. H. (2002). The neural basis of human error processing: reinforcement learning, dopamine, and the error-related negativity. Psychological Review, 109(4), 679–709. doi:10.1037/0033-295X.109.4.679

Holroyd, C. B., & Umemoto, A. (2016). The research domain criteria framework: The case for anterior cingulate cortex. Neuroscience and Biobehavioral Reviews, 71, 418–443. doi:10.1016/j.neubiorev.2016.09.021

Huang, J., Shum, D. H. K., & Chan, R. C. K. (2013). Executive inhibition: A study of postcommission error slowing and postomission error speeding. PsyCh journal, 2(3), 161–166. doi:10.1002/pchj.25

Iacobucci, D., Posavac, S. S., Kardes, F. R., Schneider, M. J., & Popovich, D. L. (2015). Toward a more nuanced understanding of the statistical properties of a median split. Journal of Consumer Psychology, 25(4), 652–665. doi:10.1016/j.jcps.2014.12.002

IBM Cor (2016). IBM SPSS Statistics for Windows, Version 24.0. Armonk, NY: IBM Corp.

Ladouceur, C. D., Dahl, R. E., & Carter, C. S. (2007). Development of action monitoring through adolescence into adulthood: ERP and source localization. Developmental Science, 10(6), 874–891. doi:10.1111/j.1467-7687.2007.00639.x

Leonard, H. C., Bernardi, M., Hill, E. L., & Henry, L. A. (2015). Executive functioning, motor difficulties, and developmental coordination disorder. Developmental Neuropsychology, 40(4), 201–215. doi:10.1080/87565641.2014.997933

Leversen, J. S. R., Haga, M., & Sigmundsson, H. (2012). From children to adults: motor performance across the life-span. Plos One, 7(6), e38830. doi:10.1371/journal.pone.0038830

Luck, S. J. (2014). An Introduction to the Event-Related Potential Technique (2, illustrated.). MIT Press.

Ludyga, S., Pühse, U., Gerber, M., & Herrmann, C. (2019). Core executive functions are selectively related to different facets of motor competence in preadolescent children. European journal of sport science, 19(3), 375–383. doi:10.1080/17461391.2018.1529826

Marchetti, R., Forte, R., Borzacchini, M., Vazou, S., Tomporowski, P. D., & Pesce, C. (2015). Physical and motor fitness, sport skills and executive function in adolescents: A moderated prediction model. Psychology, 06(14), 1915–1929. doi:10.4236/psych.2015.614189

McIntyre, F., Parker, H., Thornton, A., Licari, M., Piek, J., Rigoli, D., & Hands, B. (2017). Assessing motor proficiency in young adults: The Bruininks Oseretsky Test-2 Short Form and the McCarron Assessment of Neuromuscular Development. Human Movement Science, 53, 55–62. doi:10.1016/j.humov.2016.10.004

Meyer, T. J., Miller, M. L., Metzger, R. L., & Borkovec, T. D. (1990). Development and validation of the Penn State Worry Questionnaire. Behaviour Research and Therapy, 28(6), 487–495. doi:10.1016/0005-7967(90)90135-6

Neurospec. Easycap System Kit. Stans, CH.

Olatunji, B. O., Schottenbauer, M. A., Rodriguez, B. F., Glass, C. R., & Arnkoff, D. B. (2007). The structure of worry: relations between positive/negative personality characteristics and the Penn State Worry Questionnaire. Journal of Anxiety Disorders, 21(4), 540–553. doi:10.1016/j.janxdis.2006.08.005

Olvet, D. M., & Hajcak, G. (2009). The stability of error-related brain activity with increasing trials. Psychophysiology, 46(5), 957–961. doi:10.1111/j.1469-8986.2009.00848.x

Pajkossy, P., Keresztes, A., & Racsmány, M. (2017). The interplay of trait worry and trait anxiety in determining episodic retrieval: The role of cognitive control. Quarterly Journal of Experimental Psychology, 70(11), 2234–2250. doi:10.1080/17470218.2016.1230142

Perri, R. L., Spinelli, D., & Di Russo, F. (2017). Missing the target: the neural processing underlying the omission error. Brain topography, 30(3), 352–363.

Plummer, P., Eskes, G., Wallace, S., Giuffrida, C., Fraas, M., Campbell, G., … American Congress of Rehabilitation Medicine Stroke Networking Group Cognition Task Force. (2013). Cognitive-motor interference during functional mobility after stroke: state of the science and implications for future research. Archives of Physical Medicine and Rehabilitation, 94(12), 2565–2574.e6. doi:10.1016/j.apmr.2013.08.002

Psychology Software Tools, Inc. (2012). E-Prime 2.0. Pittsburgh, PA.

Querne, L., Berquin, P., Vernier-Hauvette, M.-P., Fall, S., Deltour, L., Meyer, M.-E., & de Marco, G. (2008). Dysfunction of the attentional brain network in children with Developmental Coordination Disorder: a fMRI study. Brain Research, 1244, 89–102. doi:10.1016/j.brainres.2008.07.066

Rigoli, D., Kane, R. T., Mancini, V., Thornton, A., Licari, M., Hands, B., … Piek, J. (2017). The relationship between motor proficiency and mental health outcomes in young adults: A test of the Environmental Stress Hypothesis. Human Movement Science, 53, 16–23. doi:10.1016/j.humov.2016.09.004

Rigoli, D., Piek, J. P., Kane, R., & Oosterlaan, J. (2012). An examination of the relationship between motor coordination and executive functions in adolescents. Developmental Medicine & Child Neurology, 54(11), 1025–1031.

Robbins, T. W. (2009). From behavior to cognition: functions of mesostriatal, mesolimbic, and mesocortical dopamine systems. In L. Iversen, S. Iversen, S. Dunnett, & A. Bjorklund (eds.), Dopamine Handbook (pp. 203–214). Oxford University Press. doi:10.1093/acprof:oso/9780195373035.003.0014

Sahlander, C., Mattsson, M., & Bejerot, S. (2008). Motor function in adults with Asperger’s disorder: a comparative study. Physiotherapy Theory and Practice, 24(2), 73–81. doi:10.1080/15368370701380843

Sartori, R. F., Valentini, N. C., & Fonseca, R. P. (2020). Executive function in children with and without developmental coordination disorder: A comparative study. Child: care, health and development, 46(3), 294–302. doi:10.1111/cch.12734

Schüller, T., Gruendler, T. O. J., Huster, R., Baldermann, J. C., Huys, D., Ullsperger, M., & Kuhn, J. (2018). Altered electrophysiological correlates of motor inhibition and performance monitoring in Tourette’s syndrome. Clinical Neurophysiology, 129(9), 1866–1872. doi:10.1016/j.clinph.2018.06.002

Shenhav, A., Botvinick, M. M., & Cohen, J. D. (2013). The expected value of control: an integrative theory of anterior cingulate cortex function. Neuron, 79(2), 217–240. doi:10.1016/j.neuron.2013.07.007

Silva, V., Campos, C., Sá, A., Cavadas, M., Pinto, J., Simões, P., … Barbosa-Rocha, N. (2017). Wii-based exercise program to improve physical fitness, motor proficiency and functional mobility in adults with Down syndrome. Journal of Intellectual Disability Research, 61(8), 755–765. doi:10.1111/jir.12384

Sollinger, A. B., Goldstein, F. C., Lah, J. J., Levey, A. I., & Factor, S. A. (2010). Mild cognitive impairment in Parkinson’s disease: subtypes and motor characteristics. Parkinsonism & Related Disorders, 16(3), 177–180. doi:10.1016/j.parkreldis.2009.11.002

Steinborn, M. B., Flehmig, H. C., Bratzke, D., & Schröter, H. (2012). Error reactivity in self-paced performance: Highly-accurate individuals exhibit largest post-error slowing. Quarterly Journal of Experimental Psychology, 65(4), 624–631. doi:10.1080/17470218.2012.660962

Stuhr, C., Hughes, C. M. L., & Stöckel, T. (2018). Task-specific and variability-driven activation of cognitive control processes during motor performance. Scientific Reports, 8(1), 10811. doi:10.1038/s41598-018-29007-3

Themanson, J. R., Rosen, P. J., Pontifex, M. B., Hillman, C. H., & McAuley, E. (2012). Alterations in error-related brain activity and post-error behavior over time. Brain and Cognition, 80(2), 257–265. doi:10.1016/j.bandc.2012.07.003

van der Fels, I. M. J., Te Wierike, S. C. M., Hartman, E., Elferink-Gemser, M. T., Smith, J., & Visscher, C. (2015). The relationship between motor skills and cognitive skills in 4-16 year old typically developing children: A systematic review. Journal of science and medicine in sport / Sports Medicine Australia, 18(6), 697–703. doi:10.1016/j.jsams.2014.09.007

Weinberg, A., Kotov, R., & Proudfit, G. H. (2015). Neural indicators of error processing in generalized anxiety disorder, obsessive-compulsive disorder, and major depressive disorder. Journal of Abnormal Psychology, 124(1), 172–185. doi:10.1037/abn0000019

Yokota, Y., Soshi, T., & Naruse, Y. (2019). Error-related negativity predicts failure in competitive dual-player video games. PloS one, 14(2), e0212483.

Yoon, D. Y., Scott, K., Hill, M. N., Levitt, N. S., & Lambert, E. V. (2006). Review of three tests of motor proficiency in children. Perceptual and Motor Skills, 102(2), 543–551. doi:10.2466/pms.102.2.543-551

